# Biological Sex Influences the Contribution of Sign-Tracking and Anxiety-Like Behaviour toward Remifentanil Self-Administration

**DOI:** 10.1101/2022.10.28.514235

**Authors:** Alicia Zumbusch, Ana Samson, Chloe Chernoff, Brandi Coslovich, Tristan Hynes

## Abstract

Most people sample addictive drugs, but use becomes disordered in only a small minority. Two important factors that influence susceptibility to addiction are individual differences in personality traits and biological sex. The influence of traits on addiction-like behaviour is well characterized in preclinical models of cocaine self-administration, but less is understood in regards to opioids. How biological sex influences trait susceptibility to opioid self-administration is likewise less studied than psychostimulants. Thus, we sought to elucidate how biological sex and several addiction-relevant traits interact with the propensity to self-administer the opioid remifentanil. We first screened female (*n*=19) and male (*n*=19) rats for four addiction-relevant traits: impulsivity, novelty place-preference, anxiety-like behaviour, and attribution of incentive value to reward cues. Rats were then trained to self-administer remifentanil in a “conflict model” of drug self-administration. Rats had to endure a mild electric shock to access the response manipulandum that triggered an intravenous infusion of remifentanil. In male rats, high anxiety-like behaviour was positively correlated with the number of drug infusions if the shock level was low or completely absent. In females, sign-tracking was predictive of greater resistance to punishment during drug seeking; an effect that was mediated by anxiety-like behaviour. Females consumed more remifentanil under all conditions, and their drug seeking persisted in the face of significantly greater current than males. These findings demonstrate that the influence of behavioural traits over the propensity to self-administer opioids is dependent upon biological sex.

## Introduction

Drug addiction is a fast-growing health epidemic in North America, costing an estimated $500 billion annually (Rehm et al., 2006; Rehm et al., 2007; Sacks et al., 2015). Addiction also has serious non-monetary costs, such as the degradation of communities, relationships, and individual mental health. Though 90% of individuals will try some form of addictive drug in their lifetime, only 12% end up having an actual addiction (Anthony et al., 1997; Gowing et al., 2015). Thus, determining the factors that render some individuals more vulnerable to develop addiction is crucial. Individual differences in personality traits play a role in addiction liability, as does biological sex (Becker et al., 2017; DuPont, 1998; Ersche et al., 2010). Concluding whether the trait is a precursor or consequence of drug use is not possible using correlational human data. However, there are several animal models designed to operationalize human personality traits which have been widely used to explore the influence of traits on addiction behaviours. The traits with the most empirical evidence to support their link to addiction are: impulsive-like behaviour, novelty preference, anxiety-like behaviour and attribution of incentive value to reward cues (Beckmann et al., 2011; Belin et al., 2016; Ersche et al., 2012; Ersche et al., 2010; Fineberg et al., 2010; Robbins et al., 2012; Saunders & Robinson, 2013).

Impulsivity is a trait that is strongly associated with drug addiction. Individuals who suffer from impulse control disorders have a higher lifetime prevalence of disordered use of all classes of addictive drugs (Levy et al., 2018; Ohlmeier et al., 2008; Verdejo-García et al., 2008). People with substance use disorders also show higher levels of impulsivity in laboratory tests and self-reports (Allen et al., 1998; Oh et al., 2021). For psychostimulants (Belin et al., 2008; Marusich & Bardo, 2009; Perry et al., 2005), alcohol (Poulos et al., 1995), and nicotine (Diergaarde et al., 2009), the association between high impulsivity and addiction-relevant behaviour translates to rodents. However, impulsivity does *not* predict opioid self-administration in male rats (Dilleen et al., 2012; McNamara et al., 2010; Schippers et al., 2012). This lack of correlation is both at odds with what is observed in humans and furthermore is yet to be studied in female rats.

The human affective state of anxiety is also associated with addiction. Indeed, there is high comorbidity between anxiety disorders and substance use disorders in humans (Conway et al., 2002; Ipser et al., 2015; Marmorstein et al., 2010). In particular, individuals with anxiety disorders are more likely to choose and use drugs with anxiolytic properties (e.g., opioids) in an aberrant fashion (Lejuez et al., 2008; Markou et al., 1998; Martins et al., 2012). These data are in line with the theory that people use drugs to “self-medicate” to quell the aversive aspects of anxiety and support the reciprocal notion that high-anxiety individuals may be less prone to use anxiogenic drugs (e.g., cocaine; (Khantzian, 1987). However, in rats, anxiety-like behaviour predicts cocaine, but not opioid self-administration (Dilleen et al., 2012; Swain et al., 2018; Walker et al., 2009). A possible explanation for this discrepancy is that individuals are more driven to “self-medicate” in anxiogenic contexts (e.g., when an electric shock is present), though this idea has yet to be addressed in a pre-clinical study.

In humans, there is substantial interindividual variability in the enjoyment of and the tendency to seek out risky, intense, and novel experiences; a trait called sensation seeking (Zuckerman, 1984, 1994). The subject effects of drugs are one such experience that sensation seekers may pursue as they tend to produce exciting, novel experiences (Dellu et al., 1996). Indeed, sensation seekers are more likely to use psychostimulants (Ersche et al., 2010; Lookatch et al., 2012; Nielsen et al., 2012), opioids (Kosten et al., 1994), nicotine (Carton et al., 1994), and alcohol (Hittner & Swickert, 2006). In rats, sensation seeking is modelled by assessing locomotor responsivity to a novel environment or the choice of a novel environment over a familiar one (i.e., novelty place-preference) (Belin et al., 2011; Piazza et al., 1989). Responsivity to novelty predicts cocaine (Piazza et al., 1989; Piazza et al., 2000), but not opioid self-administration (Chang et al., 2022; Swain et al., 2018; Swain et al., 2020). Similarly, rats with high novelty place-preference show greater cocaine self-administration than those who are less novelty preferring (Belin et al., 2011; Belin et al., 2008). The relationship between novelty preference and opioid self-administration remains unexplored.

The cues (i.e., conditioned stimuli) associated with drugs (e.g., a syringe) have robust motivational control over drug seeking (Stewart et al., 1984). Critically, individual differences exist in the degree to which such cues rouse motivational urges. Individuals that attribute more motivational value to conditioned stimuli are referred to as sign-trackers (STs) and those who attribute comparatively less are referred to as goal-trackers (GTs) (Robinson & Flagel, 2009). Sign-trackers are more prone to exhibiting addiction-like behaviour in studies examining psychostimulants (Everett et al., 2020; Flagel et al., 2009; Saunders & Robinson, 2010; Saunders & Robinson, 2011; Saunders et al., 2013b; Tunstall & Kearns, 2015). This may be because of their aberrant attraction to cues or the constellation of other addiction-relevant traits associated with sign-tracking (Beckmann et al., 2011; Campus et al., 2019; Tomie, Aguado, et al., 1998; Tomie, Cunha, et al., 1998), Interestingly, sign-tracking does not appear to increase the vulnerability to behaviours related to opioid addiction such as drug seeking or reinstatement (Chang et al., 2022; Martin et al., 2022b). There is, however, evidence that STs are more resistant to punishment in a cocaine-seeking paradigm (Pohořalá et al., 2021).Thus, whether STs are resistant to punishment in the context of opioid self-administration is an important new area of research to pursue.

Thus far, we’ve summarized clear evidence that impulsivity, anxiety, novelty-preference/reactivity, and high attribution of incentive value to reward cues are risk factors in humans developing an addiction. Additionally, we summarized studies showing that several preclinical behavioural analogues of these traits relate to many addiction-relevant behaviours in rodent models. Nonetheless, how these same traits affects the self-administration of opioids is much less well characterized (Swain et al., 2021). This is especially the case in understanding the role of sex in the trait-addiction relationship. Given that females differ substantially from males in their patterns of opioid self-administration, this is an essential area of inquiry (Carroll et al., 2002; George et al., 2021; Klein et al., 1997; Lynch & Carroll, 1999; Thorpe et al., 2020).

In the present study, we aimed to address the outstanding gaps in research regarding the how biological sex and four addiction-related traits confer increased addiction vulnerability. Here, we screened female and male rats for impulsivity, anxiety, novelty preference, and the attribution of incentive value to reward cues. We then trained them to self-administer remifentanil in a “conflict” model (Cooper et al., 2007), where an aversive electric shock is used to model the negative consequences of drug seeking. Over sessions, the intensity of the electric shock was increased until rats abstain from drug seeking entirely allowing us to assess individual differences in resistance to punishment. The conflict paradigm also allowed us to assess differences in the acquisition of remifentanil self-administration and the relapse-like behaviour following shock-induced abstinence. We then performed correlational analyses to determine the relationship between these addiction relevant outcomes and the four traits as well as biological sex.

Based on several human correlational studies showing that impulse control impacts addiction in females more than it does in males (for review, see (Fattore & Melis, 2016), we hypothesize that high impulsivity may be a predictor of potentiated opioid self-administration in females, compared to males. Anxious individuals (i.e., those high in *trait* anxiety) may become more anxious in anxiety-provoking situations (i.e., state anxiety) making them more likely to “self-medicate” with anxiolytic/sedating drugs such as opioids. As such, we predict that in the presence of an anxiogenic electric shock, anxiety-like behaviour will positively correlate with remifentanil intake. Others have elegantly summarized evidence that the association between novelty preference and addiction-like behaviour is tenuous (O’Connor et al., 2022), so here we make no *a priori* hypothesis regarding this trait. Lastly, given that sign-trackers are resistant to punishment in cocaine-seeking paradigms (Pohořalá et al., 2021), we predict they will be likewise resistant to punishment our opioid self-administration paradigm.

## Methods

### Animals

All experiments were conducted with male (n = 19; ∼300-400g) and female (n = 19; ∼200-275g) Sprague-Dawley rats (Charles River, Saint-Constant, Quebec), between 10 and 20 weeks of age. All rats were housed under a 12:12 reverse light-dark cycle. The temperature of the room was maintained between 22 °C and 24 °C and the humidity was kept constant ∼ 30%). Rats had *ad libitum* access to water and standard chow (LabDiet® 5001, LabDiet, St. Louis, MO, USA). Procedures were executed under the approval of the Life and Environmental Sciences Animal Care Committee of the University of Calgary and adhere to the Canadian Council of Animal Care guidelines.

### Apparatus

#### Operant apparatus

Differential reinforcement of low rates of responding (DRL-10s), Pavlovian conditioned approach (PCA), remifentanil self-administration, and vocalization threshold procedures were conducted in 20 modular testing chambers (D:24 cm × W:30 cm × H:30 cm; all accessories from Med Associates Inc., St. Albans, VT, USA). Chambers were housed in sound-attenuating cubicles that had ventilation fans to generate masking noise. Med Associates software controlled the chambers and recorded all session data. For DRL-10s testing, the chambers were outfitted with a white house light, centrally located at the top of one wall, a food cup (magazine), centrally located on the opposite wall (2.5 cm above the stainless-steel mouse grid floor), and a response wheel (7.5 cm diameter; 5 cm width) flanking the magazine (left or right). For PCA testing, the chambers were outfitted with a red house light, centrally located at the top of one wall, and a magazine on the opposite wall. Magazine entries were recorded by the breaking of an infrared photo-beam. Lastly, a retractable, back-illuminated (by white LED) lever (5 cm long; 6 cm above the grid floor) was located either to the left or right of the food magazine (counterbalanced across rats). For remifentanil self-administration, operant chambers were equipped with two nosepoke response ports (2.5cm dia.) containing an LED cue light; one on the left and one on the right side of the front wall situated 2cm above the grid floor. Entries into these nosepoke ports were logged via an internal infrared photo-sensor. These operant chambers were also equipped with a cantilevered drug delivery arm and vascular access tether system (PMH-110-SAI, Med Associates Inc., St. Albans, VT, USA). Outside the sound-attenuating cubicle, a 10mL syringe containing the remifentanil solution was loaded into a variable infusion rate syringe pump (PMH-107, Med Associates Inc., St. Albans, VT, USA) and connected to the vascular access system via polyethylene tubing. The pump was calibrated to deliver a 50uL infusion over 3.7 s (i.e., 13.5 uL/s). Using a custom wiring harness (ENV-406AM, Med Associates Inc., St. Albans, VT, USA) connected to a variable shock generator (PMH-401B, Med Associates Inc., St. Albans, VT, USA), two thirds (W:20cm x D:24cm) of the grid floor in front of the nosepoke ports was electrified. The rear W:10cm x D:24cm of the floor remained un-electrified. In this arrangement, the rat was required to cross the electrified section to gain access to the response ports, but could remain safe in the rear of the chamber (Cooper et al., 2007). Vocalization thresholds were tested in operant chambers wherein the entire floor was electrified, such that the mild foot shock was inescapable. For individual rats, DRL-10, PCA, remifentanil self-administration, and vocalization threshold were conducted in different operant chambers.

#### Open-Field Apparatus

The open-field arena was an open-top square container (D: 120cm x W: 120cm; H: 45cm) constructed from black open-cell PVC foam with a matte finish (1cm thick). The apparatus was housed in a low light (30lux), dedicated test room, and was situated between two white noise generators (Marpac Dohm-DS All Natural Sound Machine, Marpac Dohm, Wilmington, NC, USA), which produced 62dB ambient noise to mask external sounds. An overhead camera (Basler Inc., Exton, PA, USA) fed video from the arena to Noldus EthoVision video-tracking software (Noldus Information Technology B.V., Wageningen, The Netherlands). Using Noldus EthoVision software features, the test arena was divided into the periphery (defined as the outer 30cm perimeter of the box) and the centre (defined as the central 60cm x 60cm area) for analyses; however, the floor apparatus featured no overt markings.

#### Novelty Place-Preference Apparatus

Rats were tested in identical novelty place-preference apparatuses made of same PVC foam described above (D: 90cm; W: 30cm; H: 45cm). Two equal-sized compartments with black and white striped walls (D: 30cm; W: 30cm; H: 45cm), were connected by a central conduit (D: 30cm; W: 10cm; H: 45cm). The stripes in the chambers were horizontal in one chamber (3.5cm wide), and vertical in the other (7.5cm wide) yielding equal amounts of black and white wall space. The connecting hallway had black walls and the floor of all compartments had no overt features. Each striped compartment was closed off by a discreet, sliding guillotine door, connecting them to the central conduit. The three place-preference apparatuses were located next to each other in a dedicated low light (30lux) test room and flanked by two white noise generators (as above). Video tracking, recording, and analyses were performed with Noldus EthoVision video-tracking software (also as above).

### Assessment of Behavioural Traits

Behavioural trait assessments began two weeks after animals had arrived in the vivarium (10 weeks of age) and lasted for ∼5 weeks (DRL-10: 18 days; PCA: 5 days; anxiety-like behaviour: 1 day, novelty place preference: 1 day; animals were not tested on weekends and holidays). The order in which individuals were assessed on each trait was randomized.

#### Impulsivity - Differential Reinforcement of Low rates of Responding (DRL)

One week before testing commenced, rats were fed approximately 10% of their body mass daily in standard rat chow to gradually reduce them to 90% of their free-feeding weight. Food restriction lasted the duration of DRL-10s testing. The day before training, approximately forty 45mg banana-flavoured sucrose pellets (BioServ, #F0059, Frenchtown, NJ, USA) were introduced to rats’ home cage to familiarize them with the foodstuff. The following day, all rats experienced magazine training where they are placed into an operant chamber and given 25 banana pellets on a variable inter-trial interval (ITI-30s) schedule into the magazine. During this session, rats learned that pellets become available inside the magazine. No manipulanda were available in the chambers for this session. We considered rats magazine trained if they consumed all the banana pellets during this session (approximately 15mins). Next, rats were trained to spin a response wheel for delivery of a banana pellet into the magazine. Each quarter turn of the wheel was considered a single response and prompted the delivery of a pellet (FR-1 schedule). Rats were considered trained if they earned 100 pellets during this session. Next, we tested rats on a DRL-10s schedule of reinforcement in 30min sessions for 18 consecutive days. In DRL-10s, rats only earned a pellet if at least 10s had elapsed since the last reinforced response. Premature responses (i.e., responses made before the waiting period had elapsed) were not rewarded and resulted the waiting period being reset. For each session, the number of responses made, and reinforcements earned were recorded. Response efficiency was calculated as a percentage of total responses that were reinforced. Over the 18-day testing period, efficiency measures were averaged over 3-day epochs to produce 6 efficiency score blocks. Impulsivity is inversely related to efficiency on the DRL-10s schedule of reinforcement. That is, rats with high efficiency scores are considered less impulsive, whereas rats with low efficiency scores are considered more impulsive.

#### Novelty preference – Novelty place preference test

The novelty place-preference procedure consisted of two phases: habituation and novelty preference testing. Rats were brought into the behavioural testing room and placed in one of the large apparatus compartments (horizontal striped walls or vertical striped walls; distribution counterbalanced across groups). Rats explored and habituated to this compartment for 30mins. At the end of this habituation phase, the central conduit alley was opened, allowing rats to explore the whole apparatus (central alley, familiar, and novel compartments) for an additional 30mins (novelty preference test phase). Noldus EthoVision software recorded distance travelled and time spent in each compartment. Novelty preference was indexed as the proportion of time spent, and distance travelled, in the novel compartment during the novelty preference test phase.

#### Anxiety-like behaviour – Open field test

We placed rats in the apparatus and allowed them to explore for 20-minutes freely. At the time of testing, rats were completely naïve to the open field apparatus. We virtually ascribed a 10cm wide boundary around the inside periphery of the arena. A greater amount of time spent in the centre of the arena (i.e., medial of the periphery) was operationalized as low anxiety-like beahviour.

#### Attribution of incentive value to reward cues (sign-tracking) – Pavlovian conditioned approach (PCA) procedure

Rats underwent five, daily sessions of Pavlovian conditioning. Each of these conditioning sessions consisted of 25 trials. Each trial entailed the presentation of a back-illuminated lever (conditioned stimulus; CS) for 8s, after which the lever was retracted and 1 banana pellet (unconditional stimulus; US) was delivered into the magazine. The 25 trials (CS-US pairings) occurred on an ITI-90s (30-150s) schedule. Sessions lasted between 35 and 50mins. Lever deflections (i.e., contacts), entries into the magazine (i.e., contacts), latency to the first lever deflection, and latency to magazine contact during lever-CS presentation were recorded. In addition, we recorded magazine contacts during the non-CS period. Note that pellet delivery occurred irrespective of the rats’ behaviour and responding.

Following completion of Pavlovian training, we calculated a composite index to quantify. The PCA index for each rat for each day is a value between -1.0 and +1.0 (with exclusive, consistent, and instantaneous magazine responding corresponding to -1.0; likewise responding on the lever-CS corresponding to +1.0). The PCA index for days 4 and 5 of the daily training sessions were then averaged to produce a single composite index (classification PCA index) for each rat that encompassed these three facets of the attribution of incentive salience to reward-predictive stimuli (for PCA calculation formulae, see Hynes et al., 2017).

### Intravenous catheterization surgery

Rats were anesthetised with a cocktail of ketamine and xylazine (Ketaset, 42.5mg/kg i.m.; Rompun, 7.5 mg/kg i.m.), and implanted with chronic intravenous catheters, exiting through an interscapular threaded port as described by Crombag et al., 2000. Singly housed, the rats were then allowed to recover in their home cages for 5 days before commencing acquisition of self-administration. To maintain patency, catheters were flushed daily with 0.20mL of heparinized saline (50 U/mL in 0.9% saline; Sandoz Canada Inc) containing gentamicin sulphate salts (10% w/v; Sigma-Aldrich) as a prophylactic antibiotic.

### Remifentanil self-administration

Remifentanil self-administration began one week after the last behavioural trait had been assessed, such that rats were ∼17 weeks of age when the assay began. Self-administration testing was conducted every day (i.e., on weekends and holidays), until rats abstained. Rats were brought into the operant room, flushed as described above, and placed in the operant chambers. In the operant chambers, the rats were tethered to the vascular access system via the interscapular threaded port.

#### Acquisition of remifentanil self-administration

No current was delivered to the grid floor during these acquisition sessions. One nosepoke was “inactive”, in which responding was inconsequential. Responding (FR1) in the “active” nosepoke resulted in the delivery a 3.0 µg/kg infusion of remifentanil hydrochloride in saline (Teva, Stouffville, ON, Canada) in a 50 µL bolus over 3.7 s and a 10 s illumination of the LED cue light. The side on which the “active” nosepoke was located was counterbalanced across operant boxes. Following each infusion, a 10 s timeout was imposed, during which responding all responding was inconsequential. Training was conducted for 9 sessions, each of which terminated when rats reached a pre-determined infusion criterion (i.e., 3 sessions of 10 infusions, 3 sessions of 20 infusions, and 3 sessions of 40 infusions). Following each session, rats were again flushed (as above) and returned to their home cages. After the 9^th^ session, patency was confirmed by loss immediate of muscle tone with the intravenous delivery of 0.20 mL of a dilute ketamine solution (10% v/v).

#### Self-administration at baseline (0.00 mA)

The day following acquisition, rats were brought into the operant room, flushed, placed into the operant chambers, and tethered to the vascular access system. As above, responding (FR1) in the active nosepoke port delivered an infusion of remifentanil along with illumination of the cue light, followed by a timeout. Again, responding in the inactive nosepoke port was inconsequential. These sessions were not terminated at infusion criteria, rather rats freely respond for infusions/cue-light for 1 hour, after which the session ended.

#### *Remifentanil self-administration under increasing conflict (0.05* m*A to 1.00* m*A)*

The day following baseline, negative consequences were imposed for remifentanil seeking/taking. To gain access to the nosepoke, the rat was required to cross an electrified portion of floor. As first described by Cooper and colleagues, an electric current was constantly imposed over the front two-thirds of the operant chamber while the back one third remained un-electrified (Cooper et al., 2007). The current was increased by increments of 0.05 mA per session until all animals had abstained. As with the baseline session, these sessions were terminated after one hour had elapsed. As the current was increased, all rats eventually abstained from responding (i.e., achieved 0 infusions in 1hr for 3 consecutive sessions), with the vast majority doing so prior to 0.450 mA. A small minority appeared to be shock-resistant, persisting until 1.000 mA in one case.

#### Reinstatement

Once individual rats had fully abstained from responding for remifentanil, they spent 7 days in their home cage without access to the drug or being placed in the operant chamber. Following the 7-day period, rats were placed in the electrified operant chamber with the shock intensity set at 50% of the specific abstinence threshold of that rat (Saunders et al., 2013a). Rats were connected to the intravenous access tethers, but the syringe pumps were loaded with 0.9% saline, as opposed to remifentanil. The nosepoke cue light was then illuminated for 20s every 3 minutes and the number of responses made in the nosepoke hole were recorded for the duration of the 30-minute reinstatement session.

### Vocalization Threshold

To control for the possibility that individual differences in pain threshold could account for some of the observations of this experiment, we obtained a control measure of pain threshold. Following self-administration, rats were placed in the operant chambers with a completely electrified grid floor, lacking any manipulanda. Starting at 0.00 mA and incrementally elevating the current by 0.05 mA every 30s, each rat was monitored until it vocalized. The shock generator was then immediately inactivated, and the current at which vocalization occurred was recorded.

## Statistical Analyses

Repeated measures ANOVAs with session (or electrification current) as set as the within-subjects factor and sex as the between-subjects factor were used in the analysis of DRL-10, PCA, acquisition of self-administration, and the low-current epoch of self-administration (i.e., 0.00 mA to 0.15 mA). One-way univariate ANOVA was used to detect the between-subjects effect of sex on open field behaviour, novelty place preference, self-administration at baseline, abstinence threshold, and vocalization threshold. Kaplan-Meier simple survival analysis was used to statistically examine sex differences in the rate at which rats abstained from responding in the high-current epoch of self-administration. For novelty preference, anxiety-like behaviour, and impulsivity; we categorized individuals as being “high” or “low” in the trait by computing a median split for each sex separately. Associations between pro-addiction traits and self-administration behaviour were analyzed via bivariate Pearson’s correlations. The mediation analysis was conducted using PROCESS Procedure for SPSS (Version 4.1; (Hayes, 2017)). Based on a PCA index ranging from -1 to 1, sign-trackers are usually categorized as those between -1 and 0.5, goal-trackers between -0.5 and -1, and intermediates between 0.5 and -0.5 (Lomanowska et al., 2011). Based on this categorization, our experimental cohort had only 1 male goal-tracker (males: 1 GT, 8 IN, 10 ST; females: 3 GT, 7 IN, 9 ST). As such, we excluded the IN category and performed a median to categorize individuals as STs or GTs.

## Results

### Sex differences in trait expression

#### Anxiety-like behaviour

As shown in Figure 1, males spent less time in the centre of the open field arena, suggesting a higher degree of anxiety-like behaviour than females (sex: F_1,36_ = 5.157, p = 0.029). Females and males did not differ in the total distance moved in the open field arena (sex: F_1,36_ = 2.186, p = 0.148) or in their average velocity while navigating the arena (sex: F_1,36_ = 0.935, p = 0.340).

**Figure 1.**
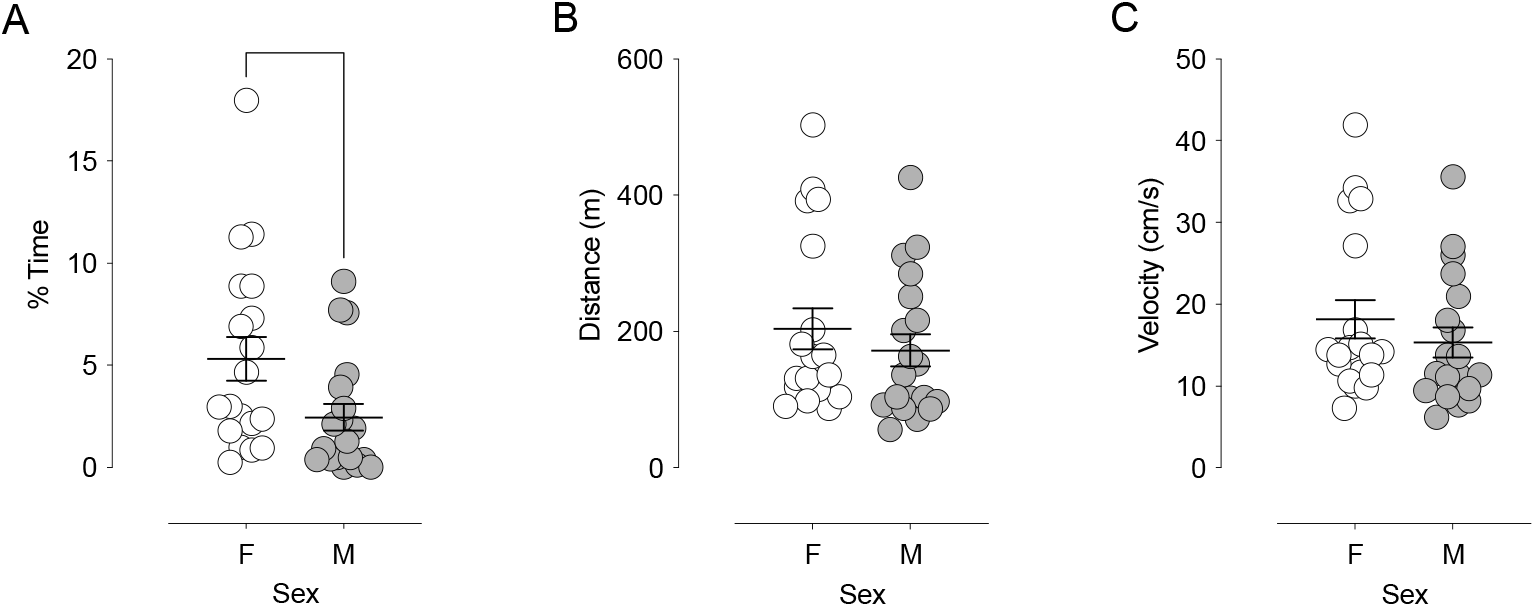
Males exhibit greater anxiety-like behaviour than females. (A) Females spent a greater proportion of time the centre of the open field arena than males. (B) Females and males did not differ in the overall distance moved or (B) average velocity in the open field arena, during the 20-minute test session. Crosshairs indicate the group mean +/- SEM. Points represent individual animals. “*” Indicates p < 0.05 between males vs. females.

#### Novelty place-preference

Indicating no difference in novelty preference, females and males did not differ in the proportion of time spent in the novel zone (sex: F_1,36_ = 1.486, p = 0.231), total distance moved, (sex: F_1,36_ = 1.491, p = 0.230) or average velocity (sex: F_1,36_ = 1.775, p = 0.191) in the novelty place-preference arena (see Supplementary Figure 1).

#### Impulsivity

Suggestive of no differences in impulsivity, in the DRL-10 task males and females did not differ from one another in responses made (session x sex: F_5,180_ = 1.685, p = 0.140); sex: F_1,36_ = 0.100, p = 0.754), reinforcers earned (session x sex: F_5,180_ = 0.395, p = 0.852); sex: F_1,36_ = 0.084, p = 0.774), or efficiency (session x sex: F_5,180_ = 0.466, p = 0.967; sex: F_1,36_ = 0.289, p = 0.594) (see Supplementary Figure 2).

#### Attribution of incentive value to reward cues

As shown in Supplementary Figure 3, females and males showed similar probabilities of approaching the magazine (session x sex: F_4,144_ = 0.933, p = 0.447; sex: F_1,36_ = 1.681, p = 0.203), number of magazine entries (session x sex: F_4,144_ = 0.748, p = 0.561; sex: F_1,36_ = 0.441, p = 0.511), and latencies to approach the magazine (session x sex: F_4,144_ = 1.199, p = 0.314; sex: F_1,36_ = 0.730, p = 0.398). Likewise, no sex differences were observed in lever-directed behaviour [probability -- (session x sex: F_4,144_ = 0.230, p = 0.921; sex: F_1,36_ = 1.150, p = 0.291); contacts -- (session x sex: F_4,144_ = 0.696, p = 0.596; sex: F_1,36_ = 0.279, p = 0.601); latency -- (session x sex: F_4,144_ = 0.448, p = 0.774; sex: F_1,36_ = 1.117, p = 0.298). PCA index also did not differ between sexes (session x sex: F_4,144_ = 0.891, p = 0.471; sex: F_1,36_ = 0.072, p = 0.790)

## Sex differences in remifentanil self-administration

### Acquisition of remifentanil self-administration

As shown in Figure 2, over the 9 acquisition sessions where the number of infusions of remifentanil was limited, the rate of remifentanil intake increased in both sexes (session: F_8,288_ = 3.46, p < 0.001; linear session: F_1,36_ = 11.823, p = 0.001). Females exhibited a higher rate of remifentanil intake overall (sex: F_1,36_ = 4.615, p = 0.039). Females also made more responses on the active nosepoke than males (sex: F_1,36_ =10.862, p = 0.001). Males and females did not differ in the number of inactive responses made (sex: F_1,36_ = 0.024, p = 0.879).

**Figure 2.**
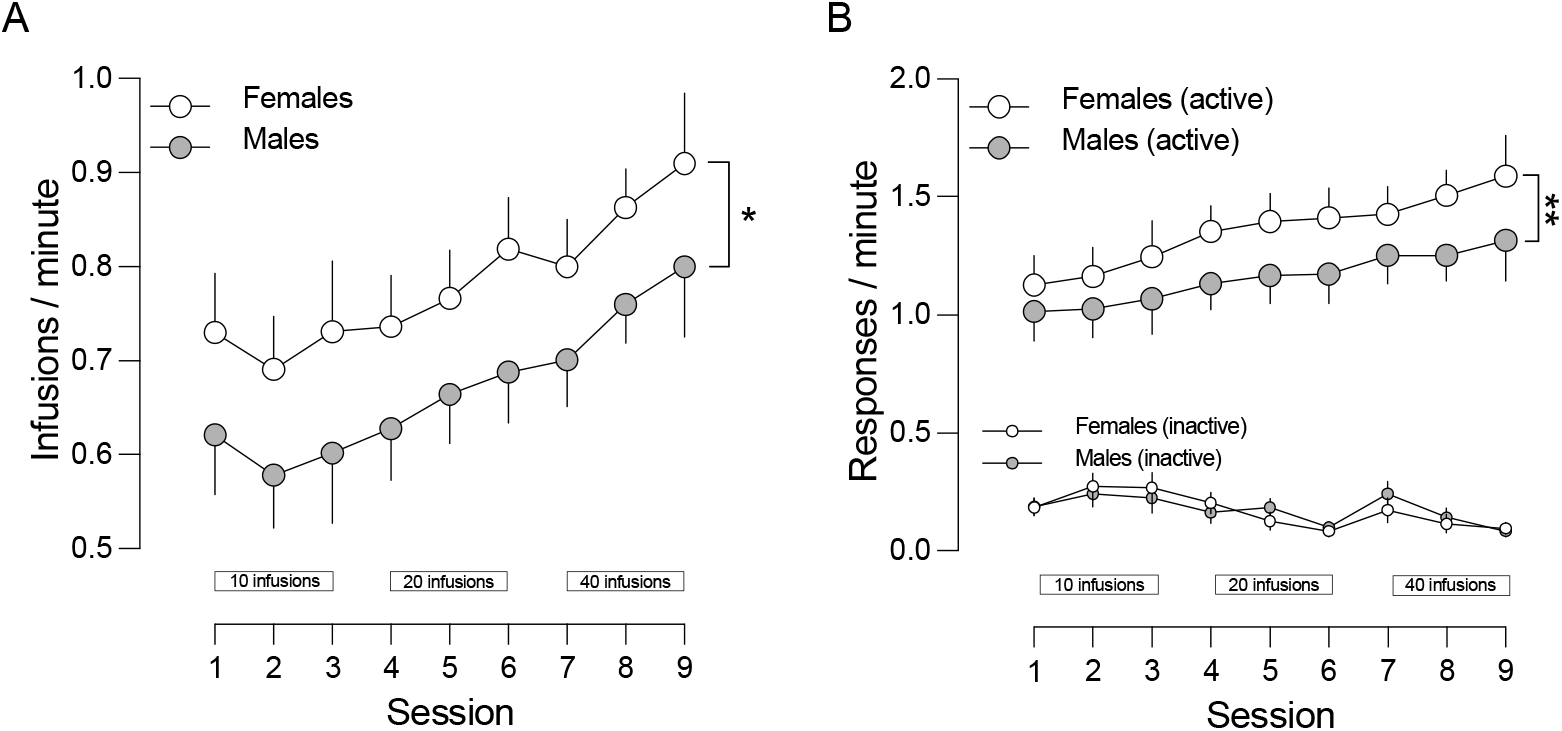
Females acquired remifentanil self-administration mor rapidly than males. (A) Females reached infusion criteria faster than males throughout all acquisition sessions. (B) Females responded in the active nosepoke more frequently than males during all phases of acquisition. Males and females did not differ in responding in the inactive nosepoke. Infusion criteria for a given session are bounded in rectangles at the bottom of the plot. Individual points represent the group mean +/- SEM. “*” Indicates p < 0.05 and “**” indicates p < 0.001 between males vs. females.

### Sex differences in remifentanil self-administration at baseline at no current conditions (0 mA to 0.15 mA)

In the first one 1-hour session where the floor was not electrified, but the number of infusions of remifentanil was not limited, females took more infusions (F_1,36_ = 7.772, p = 0.008) and made more active responses for the drug than males (F_1,36_ = 4.805, p = 0.035). The sexes did not differ in the number of inactive responses made (F_1,36_ = 1.680, p = 0.203). For the first three increments of barrier electrification (i.e., from 0.05 mA to 0.15 mA), females took more infusions (sex: F_1,36_ = 12.971, p < 0.001) and made more responses for remifentanil than males (sex: F_1,36_ = 4.102, p = 0.05). The sexes did not differ in the number of inactive responses made during the first three increments of barrier electrification (F_1,36_ = 0.957, p = 0.334) (see Figure 3).

**Figure 3.**
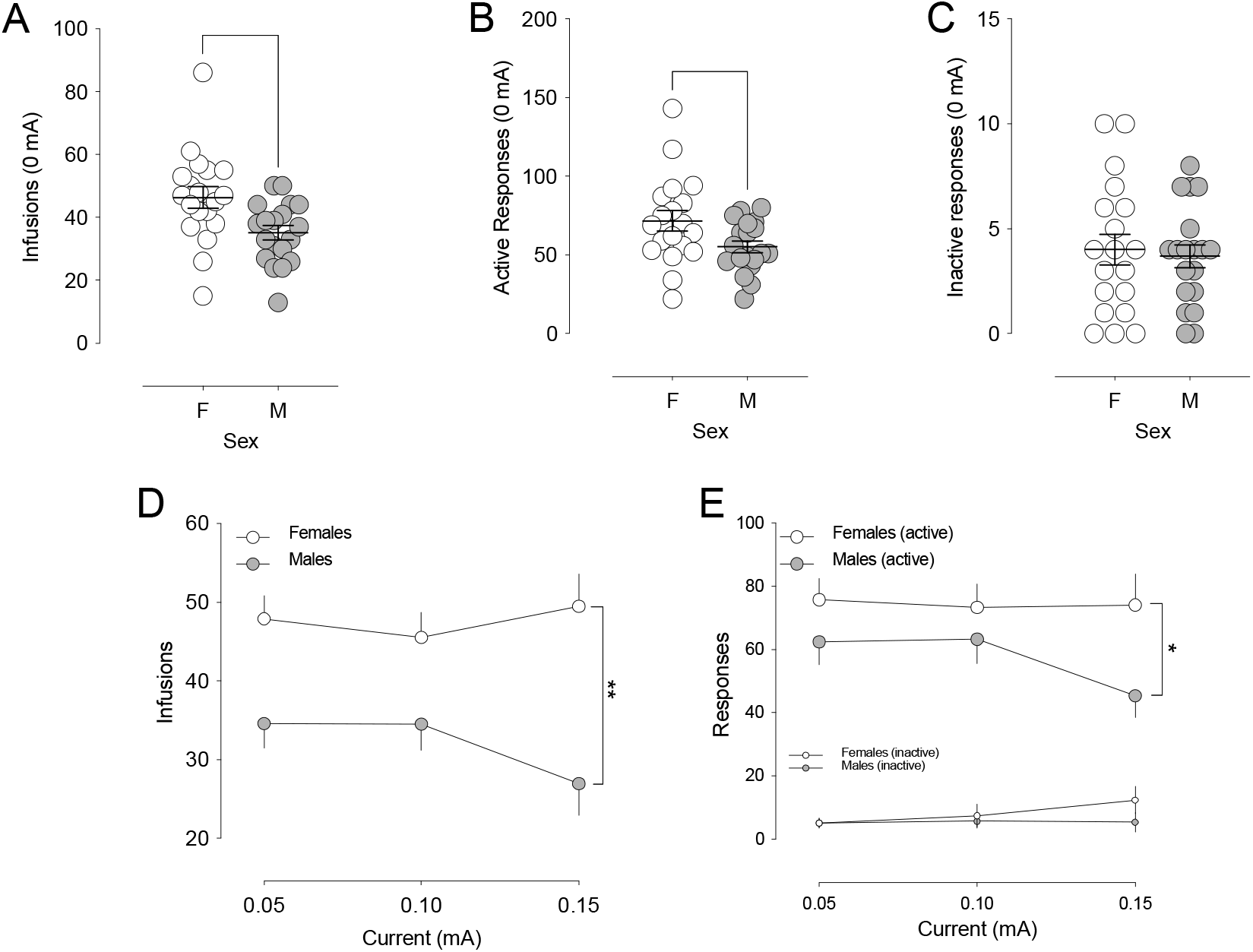
Females self-administer more and respond more for remifentanil than males under no/low current conditions. (A) When the floor is not electrified Females achieve more infusions of remifentanil than males and (B) make more responses in the active nosepoke during the 1-hour test sessions. (C) Males and females do not differ in responding in the inactive nosepoke. Crosshairs indicate the group mean +/- SEM. Points represent individual animals. (D) When the floor was electrified at low levels, females achieved more infusions of remifentanil than males and (E) made more responses in the active nosepoke hole. Females and males did not differ in responding in the inactive nosepoke. Points represent the group mean +/- SEM. “*” Indicates p < 0.05 and “**” indicates p < 0.001 between males vs. females.

### Sex differences in remifentanil self-administration under high current conditions (0.20 mA to 1.00 mA) and reinstatement following abstinence

As illustrated in Figure 4, At the electrification intensity of 0.2 mA, both males and females began to abstain from responding for the drug, which resulted in missing values that made repeated measures analysis inappropriate. We therefore employed Kaplan-Meier simple survival analysis to show that males abstained more readily as the electrification intensity was increased (X^2^ = 3.846, p = 0.0499). As evidenced by higher abstinence thresholds, females tolerated a greater intensity of shock to obtain remifentanil (sex: F_1,36_ = 4.158, p = 0.049). We observed no sex differences in reinstatement responding (sex: F_1,36_ = 0.257, p = 0.615). Because many animals did not respond during the reinstatement session (n_female_ = 10, n_male_ = 6), we also performed the analysis with non-responders excluded, yet still, females and males did not differ from one another (F_1,14_ = 1.223, p = 0.151).

**Figure 4.**
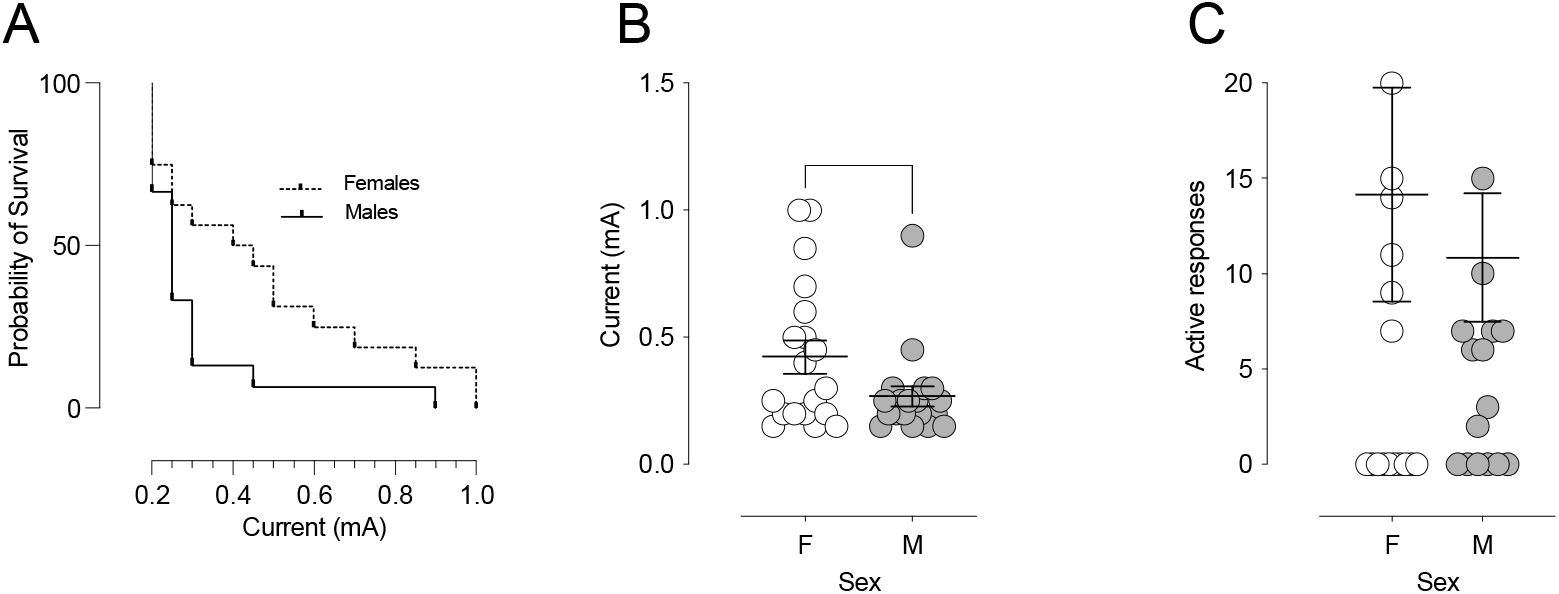
Females tolerate higher levels of shock to obtain remifentanil than males but do not differ in reinstatement following voluntary abstinence. (A) As the intensity of current was increased, males abstained from responding for remifentanil more readily than females. Survival plot depicts the cumulative percentage of rats of either sex making at least 1 response for remifentanil at a given current. (B) Females showed higher abstinence thresholds (i.e., stopped responding for remifentanil at greater currents) than males. (C) When tested one week following voluntary abstinence, females and males did not differ in the number of responses made in the active nosepoke hole during the 1-hour reinstatement session where remifentanil was not available. Individual points represent individual animals of either sex. Crosshairs indicate the group mean +/- SEM. “*” Indicates p < 0.05 and between males vs. females. We did not video record or systematically account for unusual or creative ways that the rat might avoid shock and still gain access to the drug (e.g., dangling from the tether). We did however occasionally hear excessive jangling of the tether and, upon observation, saw that some of the rats were attempting to leap toward the nosepokes.

#### Vocalization threshold

Sex had no impact on vocalization threshold (sex: F_1,36_ = 0.718, p = 0.402) (see Supplementary Figure 4), nor was any self-administration variable associated with abstinence threshold (see Supplementary Table 2), suggesting that a higher propensity to take remifentanil in the face of increasing aversive consequence was not influenced by individual differences in pain threshold. Vocalization threshold was not associated with any trait variable (see Supplementary Table 1).

#### Association of behavioural traits with remifentanil self-administration

##### Anxiety-like behaviour

Because males and females differed in both anxiety- and addiction-like behaviour, we examined the effect of anxiety-like behaviour on remifentanil self-administration in males and females separately. In neither males (session x anxiety: F_8,136_ = 0.728, p = 0.667; anxiety: F_1,17_ = 1.509, p = 0.236) nor females (session x anxiety: F_8,136_ = 0.479, p = 0.870; anxiety: F_1,17_ = 2.587, p = 0.126) did the high anxiety-like phenotype differ from the low anxiety-like phenotype in the acquisition of remifentanil self-administration. Figure 5 illustrates that males high in anxiety-like behaviours took more remifentanil than those with low anxiety-like behaviour at low current but dropped to levels similar males that were low in anxiety-like behaviour when the current reached 0.15 mA (i.e., the current after which males began to abstain) (session x anxiety: F_3,51_ = 4.225, p = 0.010; anxiety: p_0.00mA_ = 0.008, p_0.05mA_ = 0.039, p_0.10mA_ = 0.014, p_0.15mA_ = 0.569). Amount of time spent in the centre of the open field was negatively correlated with remifentanil intake during the low current sessions (r_19 @ 0.00mA_ = -0.491, p = 0.033, r_19 @ 0.05mA_ = -0.487, p = 0.034, r_19 @ 0.10mA_ = -0.596, p = 0.007, r_19 @ 0.15mA_ = -0.294, p = 0.222). We detected no such differences in females (session x anxiety: F_3,51_ = 0.305, p = 0.822). In neither sex was abstinence threshold (males - F_1,18_ = 1.093, p = 0.311; females - F_1,18_ = 0.003, p = 0.995) or reinstatement responding (males - F_1,18_ = 0.421, p = 0.681; females - F_1,18_ = 0.052, p = 0.823) affected by anxiety-like behaviour.

**Figure 5.**
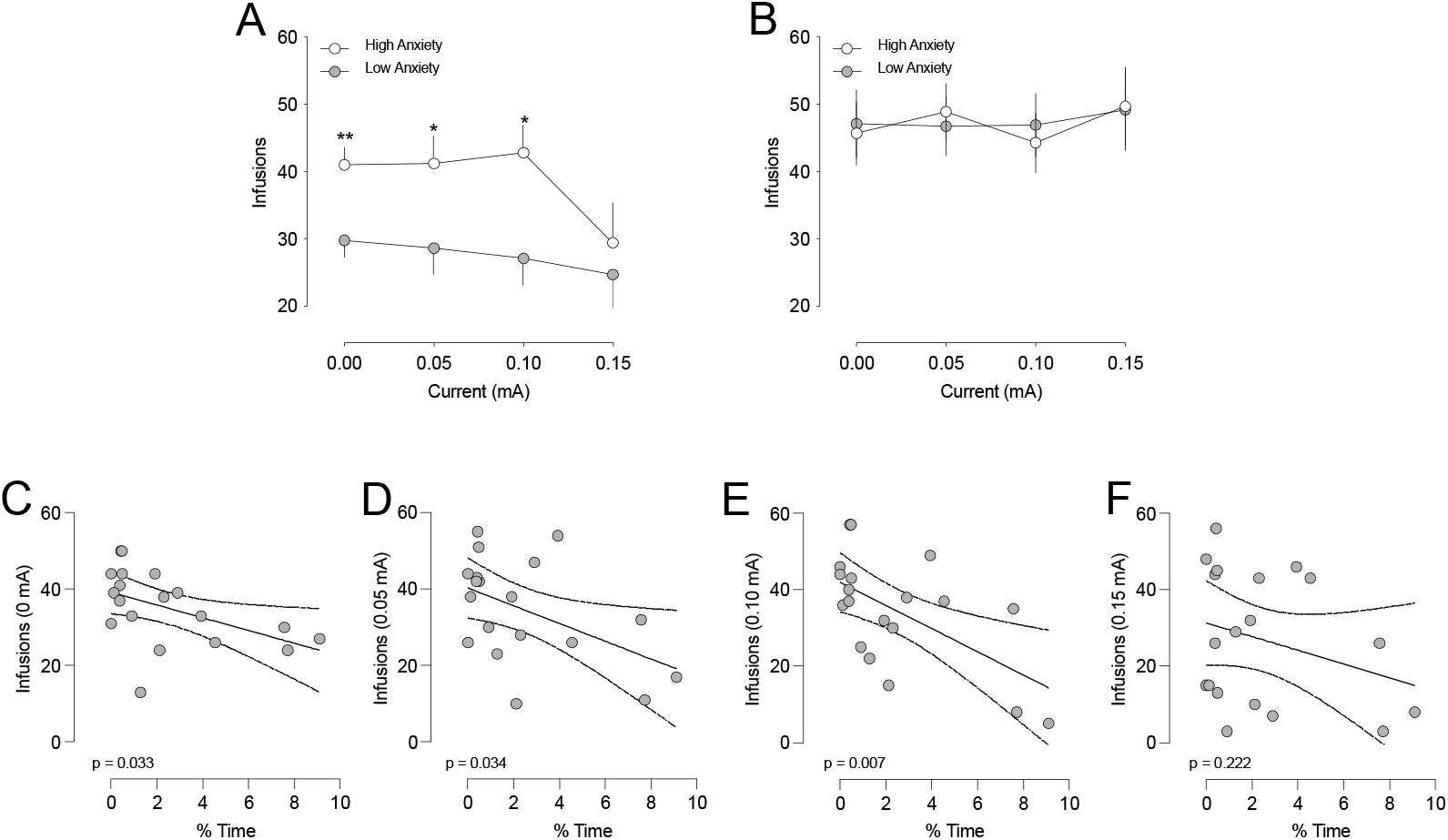
High anxiety-like behaviour predicts heightened remifentanil self-administration under no/low current conditions in females but not in males. (A) At shock levels ranging from 0.00 mA to 0.10 mA, males that exhibited a high degree of anxiety-like behaviour took more remifentanil than males exhibiting low anxiety-like behaviour; this relationship was not observed 0.15 mA. (B) Anxiety-like behaviour does not affect remifentanil self-administration in females at any shock level ranging from 0.00 mA to 0.15 mA. Points represent the group mean +/- SEM. “*” Indicates p < 0.05 and “**” indicates p < 0.001 between males exhibiting high or low anxiety-like behaviour. (C) In males, time spent in the centre of the open field was negatively correlated with the number of infusions of remifentanil achieved at 0.00 mA, (D) 0.05 mA, and (E) 0.10 mA. (F) No relationship between open field behaviour and remifentanil self was observed at 0.15 mA. P-values indicate the outcome of a 2-tailed Pearson correlation. Individual points represent individual animals. Regression line is depicted as a solid line, bound by dashed lines representing the 95% confidence intervals.

**Figure 6.**
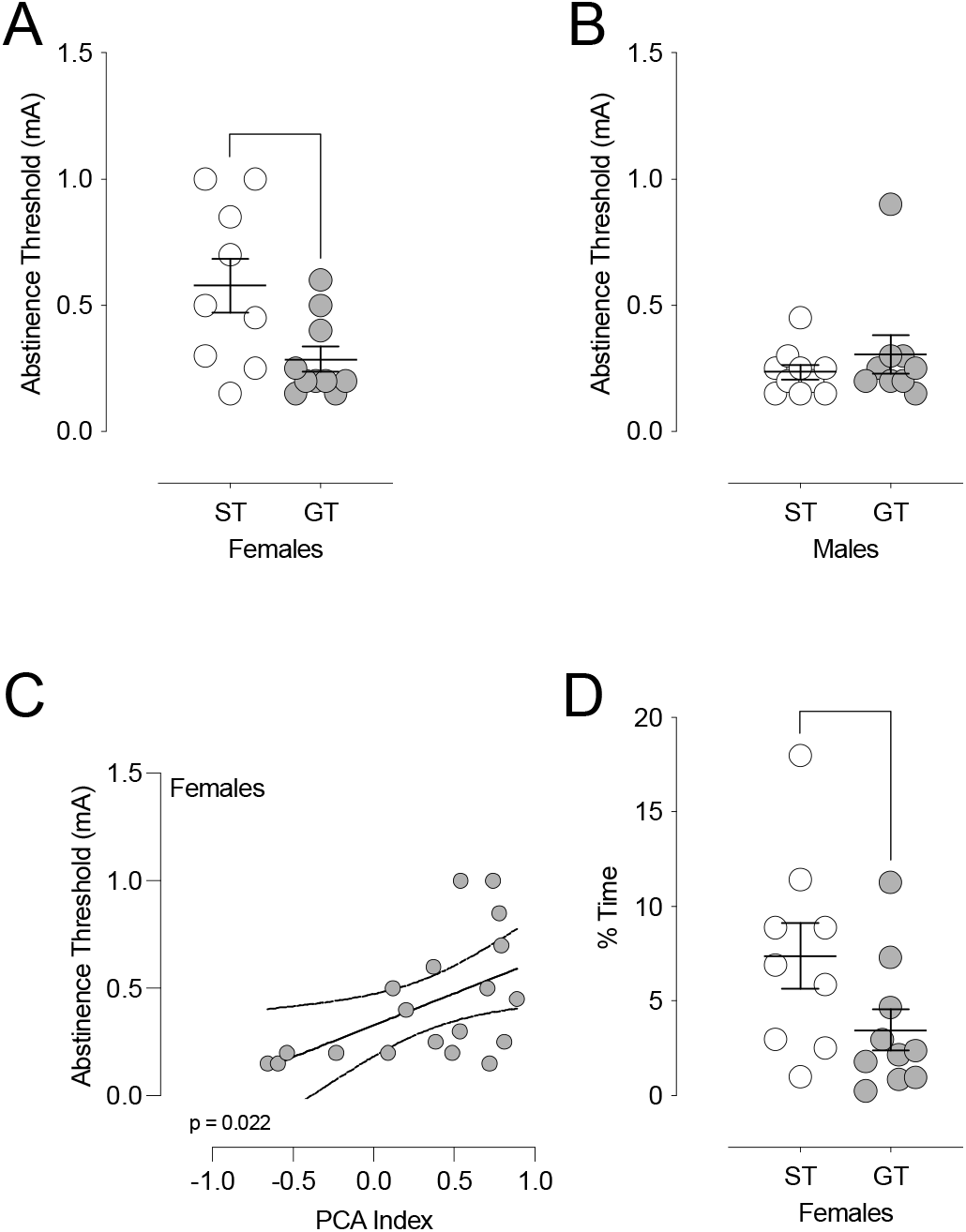
Females sign-trackers are more resistant to punishment during remifentanil self-administration – mediation by anxiety-like behaviour. (A) In females, sign-trackers (ST) exhibited higher abstinence thresholds than goal-trackers (GT). (B) In males, abstinence thresholds did not differ between STs and GTs. Crosshairs represent the group mean +/- SEM. Individual points represent individual animals. (C) A higher PCA index (i.e., greater sign-tracking behaviour) was associated with higher abstinence thresholds in females. (D) Female GTs spent a smaller proportion of time in the centre of the open field than STs, suggesting showed a greater degree of anxiety-like behaviour. “*” Indicates p < 0.05 between female STs and GTs. P-values indicate the outcome of a 2-tailed Pearson correlation. Individual points represent individual animals. Regression line is depicted as a solid line, bound by dashed lines representing the 95% confidence intervals.

##### Novelty place-preference

As depicted in Supplementary Figure 1, we detected no relationship between novelty preference and the acquisition of remifentanil self-administration (females - session x novelty: F_8,136_ = 0.592, p = 0.783; novelty: F_1,17_ = 0.969, p = 0.339; males - session x novelty: F_8,136_ = 0.582, p = 0.793; novelty: F_1,17_ = 1.550, p = 0.230) or low current self-administration (females - session x novelty: F_3,51_ = 2.648, p = 0.059; novelty: F_1,17_ = 0.000, p = 0.989; males - session x novelty: F_3,51_ = 1.571, p = 0.223; novelty: F_1,17_ = 2.501, p = 0.132). Abstinence thresholds (males - F_1,18_ = 2.759, p = 0.115; females - F_1,18_ = 0.374, p = 0.549) and reinstatement responding (males - F_1,18_ = 0.000, p = 0.990; females - F_1,18_ = 0.002, p = 0.963) were likewise unaffected by novelty preference.

##### Impulsivity

Impulsivity did not predict the acquisition of remifentanil self-administration (females - session x impulsivity: F_8,136_ = 0.441, p = 0.895; impulsivity: F_1,17_ = 0.030, p = 0.865; males - session x impulsivity: F_8,136_ = 1.013, p = 0.429; impulsivity: F_1,17_ = 0.325, p = 0.576), low current self-administration (females - session x impulsivity: F_3,51_ = 0.506, p = 0.680; impulsivity: F_1,17_ = 1.127, p = 0.303; males - session x impulsivity: F3,51 = 1.465, p = 0.235; impulsivity: F_1,17_ = 0.143, p = 0.710), abstinence thresholds (males - F_1,18_ = 1.656, p = 0.215; females - F_1,18_ = 1.555, p = 0.229), or reinstatement responding (males - F_1,18_)= 0.122, p = 0.731; females - F_1,18_ = 1.557, p = 0.229) (see Supplementary Figure 2).

##### Attribution of incentive value to reward cues

Regardless of sex, sign-trackers and goal-trackers did not differ in the acquisition of remifentanil self-administration (females - session x tracking: F_8,136_ = 1.095, p = 0.366; tracking: F_1,17_ = 0.072, p = 0.931; males - session x tracking: F_8,136_ = 0.640, p = 0.846; tracking: F_1,17_ = 0.990, p = 0.393), nor did either sex consume more remifentanil in the low current sessions (females - session x tracking: F_3,51_ = 0.010, p = 0.921; tracking: F_1,17_ = 0.046, p = 0.833; males - session x tracking: F_3,51_ = 0.611, p = 0.445; tracking: F_1,17_ = 2.465, p = 0.135). Sign-tracking females (F_1,18_ = 6.507, p = 0.021), but not males (F_1,18_ = 0.814, p = 0.380) exhibited higher abstinence thresholds. A higher PCA index was furthermore positively correlated with higher abstinence thresholds in females (r_19_ = 0.523, p = 0.022) but not in males (r_19_ = - 0.213, p = 0.382). Sign-trackers and goal-trackers did not significantly differ in terms of reinstatement responding (males-F_1,17_ = 0.053, p = 0.820; females - F_1,17_ = 2.685, p = 0.120). Female STs spent a greater proportion of time in the centre of the open field than female GTs (t_19_ = 1.948, p = 0.034), whereas the phenotypes did not differ in males (t_19_ = 0.545, p = 0.297). Though we detected a categorical difference in females, there was only a trend correlation between PCA index and abstinence thresholds in females (r = 0.355, p = 0.063; Supplementary Table 1). When time in the centre of the open field was used as a covariate in the analysis depicted in Figure 5A, STs and GTs no longer differed in abstinence thresholds (F_1,18_ = 3.485, p = 0.080). A formal using the PROCESS procedure (Hayes, 2017) showed that anxiety-like behaviour mediates the difference in abstinence thresholds between female STs and GTs (F_3,15_ = 3.366, p = 0.047).

## Discussion

We asked how biological sex, and four addiction-relevant traits interact to confer vulnerability to remifentanil self-administration under conflict. We found that females acquired remifentanil self-administration more rapidly than males, consumed more of the drug when shock levels were low, and were more resistant to shock-induced abstinence at high levels of electrical current. Males exhibited a higher degree of anxiety-like behaviour than females. There were no other sex differences in the traits assessed here. In males only, high levels of anxiety-like behaviour were associated with greater remifentanil intake at low shock levels but did not predict self-administration of the drug at high levels of shock. Sign-tracking females withstood higher levels of punishment to obtain remifentanil, and this effect was mediated by lower levels of anxiety-like behaviour in female sign-trackers.

Our finding that females acquired remifentanil self-administration more readily and did more infusions under low/no shock FR1 conditions than males is consistent with what others have reported in both rats (Thorpe et al., 2020) and mice (Anderson et al., 2021). Others have shown accelerated acquisition of self-administration in females with other opioids such as heroin (Carroll et al., 2002; George et al., 2021; Lynch & Carroll, 1999) and fentanyl (Klein et al., 1997). The evidence for sex differences in the self-administration of oxycodone in mixed, with two studies showing potentiated conflict-free FR1 responding in females (Kimbrough et al., 2020; Mavrikaki et al., 2017) and one reporting no difference (Fredriksson et al., 2020). Further studies have examined oxycodone self-administration in both sexes, but did not use sex as a factor in their analyses (Fredriksson et al., 2021). Despite the ambiguity surrounding oxycodone, most studies support the notion that female rats exhibit a heightened propensity to self-administer most opioids under a low/now shock FR1 schedule of reinforcement. There may be organizational sex differences in the central expression of mu-opioid receptors that may underlie this behavioural observation.

The rewarding effects of opioids are partly due to their effect on dopamine neurotransmission. Through both direct and indirect mechanisms, mu-opioid agonists increase the activity of dopamine release from the dopaminergic neurons of the ventral tegmental area (for review see (Fields & Margolis, 2015). Compared to males, females exhibit a lower density of midbrain mu-opioid receptors (Morley-Fletcher et al., 2003). This sex difference in mu-opioid receptor density suggests that mu-opioid agonist-induced stimulation of ventral tegmental area (VTA) dopamine neurons may be greater in males, simply by virtue of higher mu-opioid receptor availability in males. If so, females would need to self-administer more drug to achieve the same rewarding effects as males. In support of this hypothesis is evidence that intra-VTA infusions of a mu-opioid receptor antagonist results in a compensatory increase in intravenous heroin self-administration (Britt & Wise, 1983).

It is also possible that circulating gonadal hormones could account for sex differences in low/no shock FR1 responding for remifentanil. Indeed, estrous phase influences acquisition and FR1 responding for remifentanil self-administration and exogenous estradiol can modulate this self-administration (Lacy et al., 2020; Sharp et al., 2021; Thorpe et al., 2020). Given these effects of estrous cycle and gonadal hormones, a limitation of this study is that we did not account for these hormonal factors. However, the rat estrous cycle is 4-5 days long and we saw persistent sex differences in acquisition and FR1 responding in the low shock condition across all 12 consecutive days. Thus, it is unlikely that the sex differences observed here were solely due gonadal hormones.

Rats that express a higher degree of opioid withdrawal tend to seek and take opioids with enhanced vigour (Ahmed et al., 2000; Carmack et al., 2019; Lenoir & Ahmed, 2007). Though we did not assess withdrawal symptoms in the present study, we must consider the possibility that the sex differences we observed here could be influenced by sex differences in the expression of opioid withdrawal. However, males express opioid withdrawal to a greater degree than females (Bobzean et al., 2019), which leads to the prediction that *males* should express enhanced opioid self-administration, not females as we report here. This evidence, taken together with the fact that withdrawal is unlikely to occur under the present short-access conditions (Ahmed et al., 2000; Vendruscolo et al., 2011), suggests withdrawal as an unlikely factor in the observations we report here.

Sign-trackers (STs) and goal-trackers (GTs) of either sex did not differ in their self-administration of remifentanil under a low/no shock FR1 schedule (reported in males by (Chang et al., 2022)). Furthermore, we did not find that STs were any more prone to cue-induced reinstatement than GTs, which (at least in males) is in contrast to the increased reinstatement in GTs seen with cocaine (Kuhn et al., 2022; Saunders & Robinson, 2011; Saunders et al., 2013b). However, our finding that tendency to attribute incentive value to reward cues is *not* associated with cue-induced reinstatement of opioid seeking is not without precedent, as other groups have published corroborating reports (Chang et al., 2022; Martin et al., 2022a). Considerable differences likely exist across the classes of drugs regarding trait vulnerabilities to relapse-like behaviour, and the propensity to ascribe motivational value to reward cues may play into other addiction-relevant behaviours (i.e., resistance to punishment).

Regardless of their pain threshold, female rats tolerated higher intensities of electrical current to obtain remifentanil. Interestingly, this sex difference in resistance to punishment was most pronounced in STs. This finding echoes a recent report that male STs are more resistant to electric shock punishment during cocaine self-administration (Pohořalá et al., 2021). Compared to GTs, STs respond more vigorously for a cocaine-paired cue when required to cross an electric barrier make a response (Saunders et al., 2013a). Though the canonical interpretation is that STs are simply more motivated by the cocaine cue, an alternative possibility is that they are willing to tolerate greater punishment for drugs and drug cues. One explanation for the difference between STs and GTs in their resistance to punishment is that GTs more readily encode the meaning of a cue that signals shock. Flagel and colleagues posit that GTs may have a greater ability or propensity to inhibit drug-seeking behaviour when a cue signals that punishment is imminent (Flagel et al., 2021). In the “conflict model” used here, however, no discrete cue is used to indicate the occurrence of shock as the grid floor is constantly electrified. Therefore, the differential ability of STs and GTs to encode the meaning of a cue cannot explain their differences in resistance to punishment in the current data.

An alternate explanation for the increased punishment resistance in GTs than STs we observed here may relate to the variability in defensive behaviours between STs and GTs. While STs and GTs do not differ in pain sensitivity, STs do initially show less conditioned freezing to a shock-paired cue, suggesting a blunted fear response (Morrow et al., 2011; Morrow et al., 2015). In human alcohol-seeking behaviour, STs report motives associated with subjective reward, whereas GTs report fear-driven motives for alcohol-seeking (e.g., alleviation of anxiety and worry) (Liu et al., 2021). The idea that STs are less fearful than GTs prompted us to perform a post-hoc analyses to examine how anxiety-like behaviour factored into the STs’ resistance to punishment. We revealed that female STs exhibited less anxiety-like behaviour than their GT counterparts, and when the anxiety-like behaviour was controlled for, STs and GTs did not differ in abstinence thresholds. A subsequent mediation analysis confirmed that anxiety-like behaviour mediates the resistance to punishment during drug-seeking in STS that we and others report (Pohořalá et al., 2021).

The demonstration by Pohořalá and collogues that STs were resistant to punishment during cocaine self-administration was shown only in males, raising the question as to why only sign-tracking females exhibited greater resistance to punishment in the present study (Pohořalá et al., 2021). A possible answer may lay in the sex differences in the opioidergic modulation of arousal circuits, which may influence the subjective experience of pain. Noxious stimuli (e.g., foot shocks) powerfully evoke activity in locus coeruleus neurons (LC) (Liu et al., 2021). The LC projects to various nodes of the pain processing system (Senba et al., 1981; Voisin et al., 2005), where LC activity exerts an inhibitory influence, dulling the perception of pain (Maeda et al., 2009). Activation of mu-opioid receptors dampens LC activity; critically, males express far more mu-opioid receptors on LC neurons and exhibit a greater degree of mu-opioid-induced reduction in LC activity than females (Guajardo et al., 2017). Remifentanil should therefore decrease LC activity in males more than in females, and consequently, females should undergo relatively larger shock-induced increases in LC activity. This increased LC activity would decrease the subjective experience of pain in females, thereby facilitating the tolerance of higher intensities of electric shock. This line of reasoning would imply that in the self-administration of non-opioid drugs, male and female STs would be similarly resistant to punishment. However, such a prediction has not yet been empirically tested.

The relationship between anxiety and substance abuse is complicated because both state and trait anxiety likely influence drug use. Patients high in “trait” anxiety and individuals with anxiety disorders have higher relapse rates following treatment for substance use disorders (Brown & Schuckit, 1988; Driessen et al., 2000; Driessen et al., 2001; Kushner et al., 2005). Additionally, an individual dually diagnosed with substance use disorder and anxiety will report taking their abused substance to cope with the tension associated with anxiety and depression (Al’Absi, 2011). These data support the notion that the relationship between opioid use and anxiety in clinical populations is driven by a desire to self-medicate (Khantzian, 1987). However, there are opposing data that show no relationship between benign preclinical models of opioid self-administration, such as acquisition, escalation, and reinstatement paradigms (Dilleen et al., 2012; Swain et al., 2018; Swain et al., 2020). One explanation that would reconcile these oppositional data is that the relationship between anxiety-like behaviour and opioid self-administration might be more evident in an anxiogenic situation (i.e., in the presence of an electric shock). Yet contrary to our prediction, we observed the relationship only under no/low current conditions. One possible explanation is the pharmacokinetic differences between remifentanil and other opioids.

The elimination half-life of remifentanil is 8-20 minutes (Glass et al., 1999), while that of morphine and heroin (considering active metabolites) is ∼3 hours (Rook et al., 2006; Stanski et al., 1978). In a clinical setting, a single dose of heroin increases subjective mood ratings for more than 3 hours (Wallenstein et al., 1990), while the subjective effects of remifentanil rapidly decline in the first 30 minutes after administration (Black et al., 1999). These differences in duration of action imply that individuals would have to consume remifentanil more frequently than other opioids to maintain the anxiolytic effect. During a single self-administration session, rats exhibiting high levels of anxiety-like behaviour could therefore quell their negative affect with relatively few infusions of heroin or morphine, compared to remifentanil. Therefore, drug infusions for self-medication would constitute a smaller proportion of total infusions for opioids with longer durations of action, causing anxiety-like behaviour to factor in less. The fact that anxiety-like behaviour only predicted self-administration in males might be because of their higher variability within the trait and is consistent with what is typically reported in rats (Imhof et al., 1993; Johnston & File, 1991).

In summary, we have presented evidence that, under conflict in an FR1 schedule of reinforcement, female rats self-administer more remifentanil than their male counterparts and will tolerate a higher level of electric shock to obtain the drug. These findings line up with other reports that females have an increased propensity for opioid self-administration (Carroll et al., 2002; George et al., 2021; Klein et al., 1997; Lynch & Carroll, 1999; Thorpe et al., 2020). Additionally, we show that anxiety-like behaviour and sign-tracking are predictive traits of opioid self-administration in a sex-specific manner as follows:

1. High anxiety-like behaviour positively correlated with the number of infusions earned only when the shock level was low or completely absent, an effect unique to males.
2. In females, sign-tracking was predictive of greater punishment resistance during drug seeking, an effect mediated by anxiety-like behaviour.
3. Females consumed significantly more remifentanil under *all* conditions, and their drug-seeking persisted in the face of significantly greater current than males.

To our knowledge, this is the first preclinical demonstration of a relationship between any pro-addictive trait and the propensity to self-administer opioids (for review see (Swain et al., 2021). The fact that the associations described above were sex-specific and specific to these domains of self-administration (i.e., in the presence and absence of a conflict) highlights how crucial it is to use translationally relevant behavioural assays and emphasizes the vital importance of using both sexes in preclinical addiction research.

## Author Contributions

TH conceptualized and designed the experiment. TH and AZ performed the surgeries. TH, AZ, and AS conducted the behavioural testing. TH and CC performed the statistical analysis and data visualization. TH, CC, and AZ wrote the manuscript. BC & AZ edited the manuscript and provided consultation on statistical analyses.

## Supplemental Materials

**Supplementary Figure 1.**
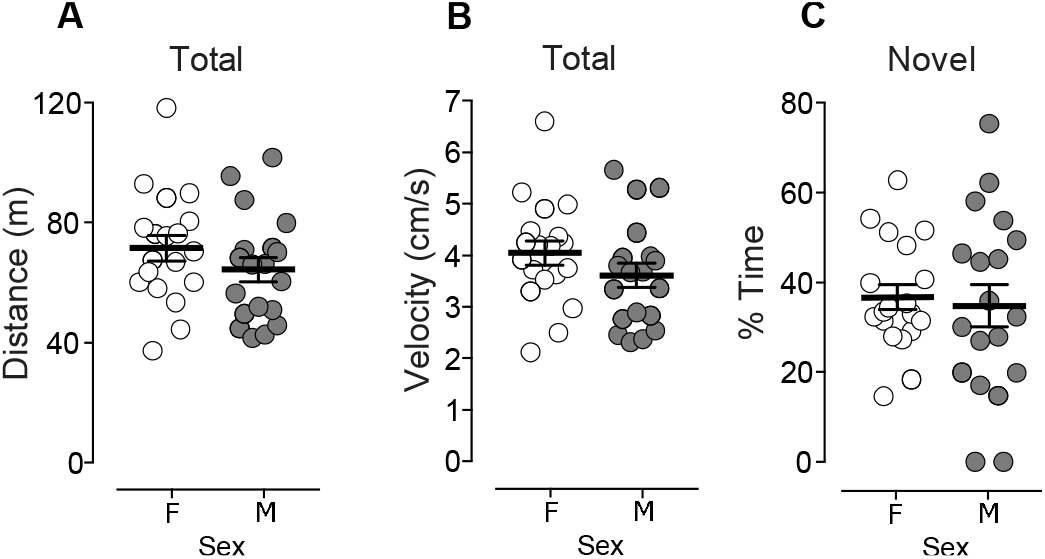
Females and males did not differ in novelty preference. (A) Females and males did not differ in the total distance moved or (B) average velocity throughout the novel place-preference arena. (C) The sexes did no differ in the proportion of time spent in the novel zone of the arena during the 20-minute test session. Individual points represent individual animals. Crosshairs indicate the group mean +/- SEM.

**Supplementary Figure 2.**
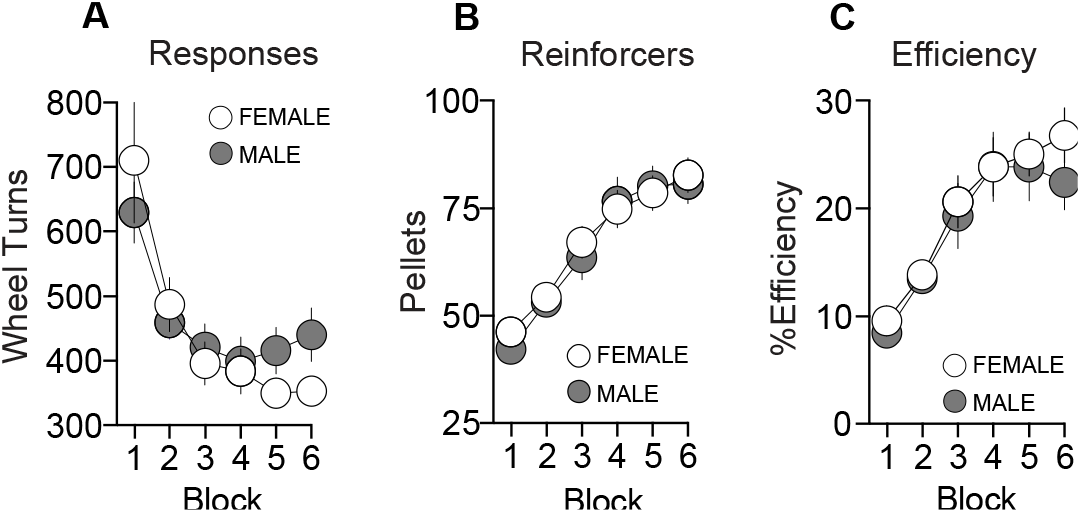
Females and males did not differ in impulsivity. (A) Males and females made a similar number of responses, (B) earned a similar number of reinforcers, (C) and exhibited similar levels of efficiency in the DRL-10 test period. Individual points represent group means +/- SEM.

**Supplementary Figure 3.**
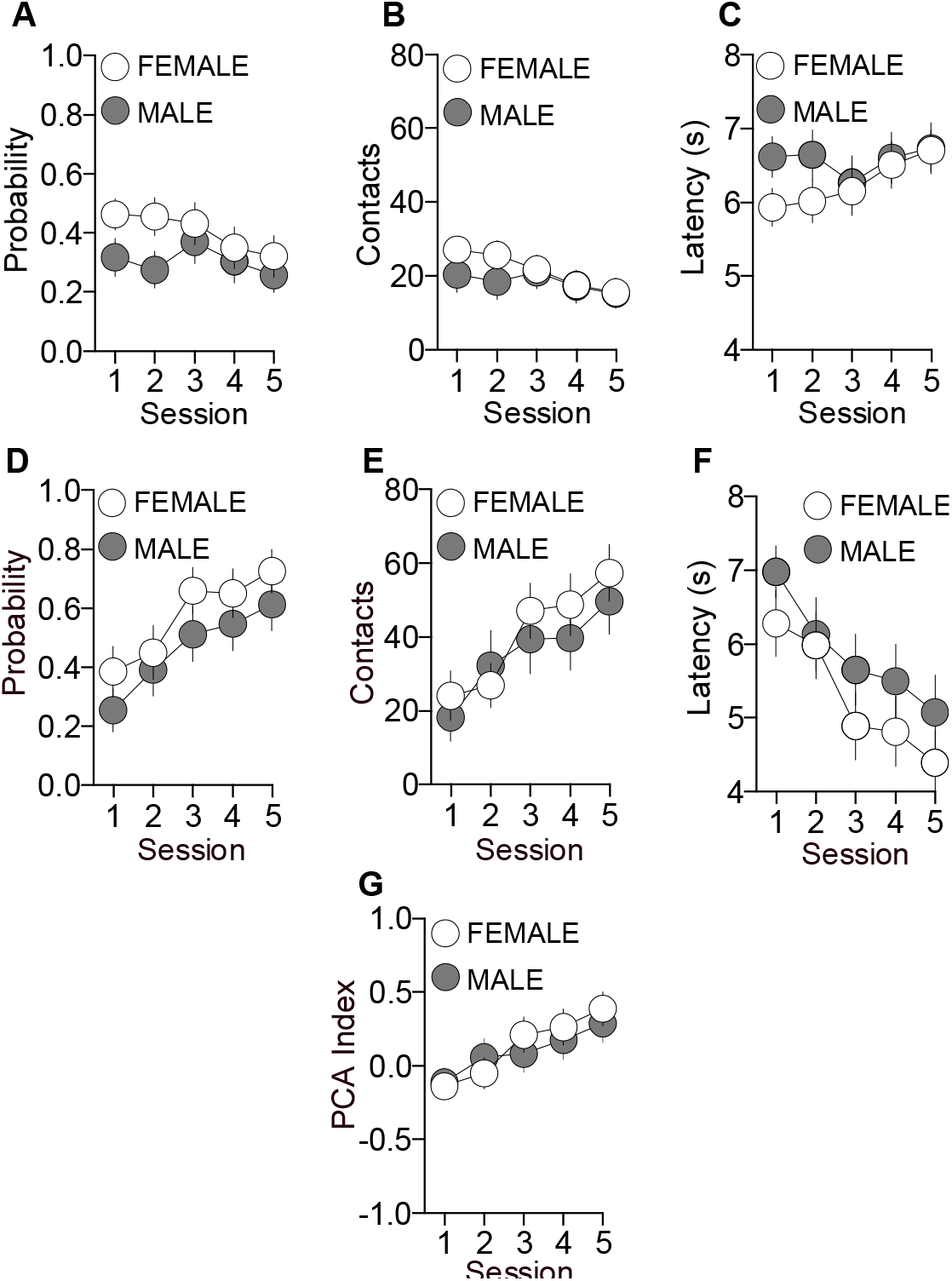
Females and males did not differ in the propensity to attribute incentive value to reward cues. (A) Males and females approached the food delivery magazine with similar probabilities, (B) made a similar number of entries into the magazine, (C) and did not differ in their relative latency to enter the magazine. (D) The sexes similarly did not differ in their (D) probabilities of contacting the lever, (E) frequency of lever contact, or (F) latency to lever contact. (G) PCA index did not differ between males and females. Individual points represent group means +/- SEM.

**Supplementary Figure 4.**
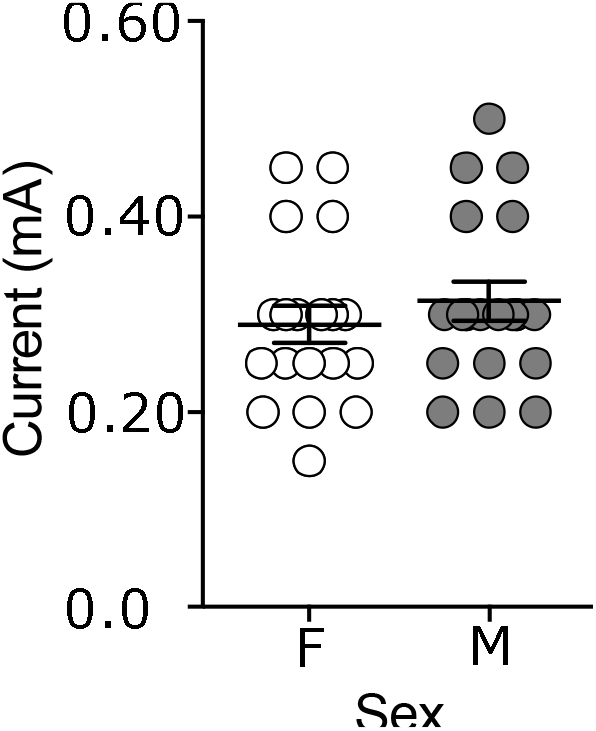
Females and males did not differ in a proxy of pain threshold. Females and males did not differ in the in intensity of electrical current require to evoke a vocalization response. Individual points represent individual animals. Crosshairs indicate the group mean +/- SEM.

**Supplementary Table 1.** Addiction-relevant traits do not correlate with one another or with vocalization threshold. In neither males nor females did traits correlate with one another or with vocalization threshold. In females a trend correlation between. Table shows Pearson correlation coefficients r. “§” indicates 2-tailed p = 0.063.

| Trait X Trait Correlations (Females) |  |  |  |  |
| --- | --- | --- | --- | --- |
|  | <u>PCA Index</u> | <u>% Time Centre OF</u> | <u>% Time Novel Zone</u> | <u>DRL Efficiency</u> |
| PCA Index | - |  |  |  |
| % Time in Centre | 0.355<sup>§</sup> | - |  |  |
| % Time in Novel | 0.063 | 0.111 | - |  |
| DRL Efficiency | 0.241 | 0.081 | -0.003 | - |
| Voc. Threshold | 0.232 | 0.057 | 0.126 | 0.419 |

| Trait X Trait Correlations (Males) |  |  |  |  |
| --- | --- | --- | --- | --- |
|  | <u>PCA Index</u> | <u>% Time Centre OF</u> | <u>% Time Novel Zone</u> | <u>DRL Efficiency</u> |
| PCA Index | - |  |  |  |
| % Time in Centre | 0.157 | - |  |  |
| % Time in Novel | 0.007 | 0.268 | - |  |
| DRL Efficiency | 0.264 | -0.049 | -0.115 | - |
| Voc. Threshold | 0.368 | 0.409 | -0.189 | -0.085 |

**Supplementary Table 3.** Sign-tracking predicts resistance to punishment in females. Low anxiety-like behaviour is associated with heightened drug intake under no/low current conditions in males. PCA index was positively correlated with abstinence thresholds in females. In males, the proportion of time spent in the centre of the open field was negatively correlated with remifentanil intake at 0.00 mA, 0.05 mA, and 0.10 mA. Table shows Pearson correlation coefficients r. Bolded values indicate significant correlations. “*” indicates 2-tailed p < 0.05. “**” indicates 2-tailed p < 0.01.

| Trait X Self-Administration Correlations (Females) |  |  |  |  |  |  |  |  |  |  |  |  |  |  |  |  |
| --- | --- | --- | --- | --- | --- | --- | --- | --- | --- | --- | --- | --- | --- | --- | --- | --- |
|  | Acquisition |  |  |  |  |  | Abstinence Threshold | Reinstatement Responses | FR1 Responding During No/Low Current Conditions |  |  |  |  |  |  |  |
|  | 10 Infusions |  | 20 Infusions |  | 40 Infusions |  |  |  | 0.00 mA |  | 0.05 mA |  | 0.10 mA |  | 0.15 mA |  |
|  | Infusions / min | Actives / min | Infusions / min | Actives / min | Infusions / min | Actives / min |  |  | Infusions | Actives | Infusions | Actives | Infusions | Actives | Infusions | Actives |
| PCA Index | -0.064 | -0.092 | 0.349 | 0.151 | 0.152 | 0.015 | 0.523* | -0.240 | -0.141 | 0.076 | 0.012 | 0.368 | -0.048 | 0.404 | -0.144 | 0.015 |
| % Time in Centre | -0.126 | 0.067 | 0.186 | 0.234 | 0.267 | 0.063 | 0.419 | -0.091 | -0.192 | -0.115 | -0.067 | 0.121 | -0.404 | -0.127 | -0.239 | -0.129 |
| % Time in Novel | 0.255 | 0.298 | -0.028 | 0.164 | 0.054 | 0.000 | -0.123 | 0.065 | 0.292 | 0.158 | 0.303 | 0.049 | 0.108 | -0.200 | -0.080 | -0.155 |
| DRL Efficiency | -0.072 | -0.132 | 0.011 | -0.111 | 0.014 | 0.130 | 0.151 | 0.105 | -0.391 | -0.280 | -0.094 | -0.036 | -0.300 | -0.180 | -0.313 | -0.088 |
| Voc. Threshold | -0.361 | -0.212 | -0.281 | -0.144 | -0.265 | 0.052 | 0.126 | -0.386 | -0.030 | 0.096 | 0.283 | 0.287 | -0.075 | 0.128 | 0.062 | 0.111 |
| Trait X Self-Administration Correlations (Males) |  |  |  |  |  |  |  |  |  |  |  |  |  |  |  |  |
|  | Acquisition |  |  |  |  |  | Abstinence Threshold | Reinstatement Responses | FR1 Responding During No/Low Current Conditions |  |  |  |  |  |  |  |
|  | 10 Infusions |  | 20 Infusions |  | 40 Infusions |  |  |  | 0.00 mA |  | 0.05 mA |  | 0.10 mA |  | 0.15 mA |  |
|  | Infusions / min | Actives / min | Infusions / min | Actives / min | Infusions / min | Actives / min |  |  | Infusions | Actives | Infusions | Actives | Infusions | Actives | Infusions | Actives |
| PCA Index | 0.362 | 0.216 | 0.298 | 0.385 | -0.005 | 0.013 | -0.213 | 0.101 | 0.075 | 0.043 | 0.377 | 0.264 | 0.248 | 0.182 | 0.397 | 0.404 |
| % Time in Centre | 0.025 | 0.109 | 0.007 | 0.052 | 0.039 | 0.084 | -0.263 | 0.164 | -0.491* | -0.295 | -0.487* | -0.221 | -0.596** | -0.306 | -0.294 | -0.183 |
| % Time in Novel | -0.358 | -0.057 | -0.234 | -0.425 | -0.028 | -0.040 | -0.263 | -0.173 | -0.225 | -0.085 | -0.171 | 0.104 | -0.310 | -0.087 | -0.057 | -0.116 |
| DRL Efficiency | 0.078 | -0.106 | 0.245 | 0.322 | 0.053 | 0.271 | 0.042 | -0.210 | -0.034 | 0.048 | 0.086 | 0.319 | 0.203 | 0.297 | 0.082 | 0.127 |
| Voc. Threshold | 0.391 | 0.057 | 0.311 | 0.087 | 0.314 | 0.249 | 0.012 | -0.002 | 0.067 | 0.111 | -0.072 | 0.145 | -0.023 | 0.099 | 0.112 | 0.200 |

